# The NSIGHT1 Randomized Controlled Trial: Rapid Whole Genome Sequencing for Accelerated Etiologic Diagnosis in Critically Ill Infants

**DOI:** 10.1101/218255

**Authors:** Josh E. Petrikin, Julie A. Cakici, Michelle M. Clark, Laurel K. Willig, Nathaly M. Sweeney, Emily G. Farrow, Carol J. Saunders, Isabelle Thiffault, Neil A. Miller, Lee Zellmer, Suzanne M. Herd, Anne M. Holmes, Serge Batalov, Narayanana Veeraraghavan, Laurie D. Smith, David P. Dimmock, Steven J. Leeder, Stephen F. Kingsmore

## Abstract

**Importance:** Genetic disorders, including congenital anomalies, are a leading cause of morbidity and mortality in infants, especially in neonatal and pediatric intensive care units (NICU and PICU). While genomic sequencing is useful for diagnosis of genetic diseases, results are usually reported too late to guide inpatient management.

**Objective:** To test the hypothesis that rapid whole genome sequencing (rWGS) increases the proportion of infants in NICUs and PICUs receiving a genetic diagnosis within 28 days.

**Design:** An investigator-initiated, partially blinded, pragmatic, randomized controlled study with enrollment from October 2014 - June 2016, and follow up until December 2016.

**Setting:** A regional neonatal and pediatric intensive care unit in a tertiary referral childrens hospital.

**Participants:** Sixty five of 129 screened families with infants aged less than four months, in neonatal and pediatric intensive care units, and with illnesses of unknown etiology, completed the study.

**Intervention:** Parent and infant trio rWGS.

**Main Outcome and Measure:** The hypothesis and end-points were formulated a priori. The primary end-point was rate of genetic diagnosis within 28 days of enrollment or first standard test order.

**Results:** Twenty six female proband infants, 37 male infants, and two infants of undetermined sex were randomized to receive rWGS plus standard tests (n=32, cases) or standard tests alone (n=33, controls). The study was terminated early due to loss of equipoise: 63% (21) controls received genomic sequencing as standard tests. Nevertheless, intention to treat analysis showed the rate of genetic diagnosis within 28 days to be higher in cases (31%, ten of 32) than controls (3%, one of 33; difference, 28% [95% CI, 10% to 46%]; p=0.003). Among infants enrolled in the first 25 days of life, the rate of neonatal diagnosis was higher in cases (32%, seven of 22) than controls (0%, zero of 23; difference, 32% [95% CI, 11% to 53%]; p=0.004). Age at diagnosis (median in cases 25 days, range 14-90 days vs median in controls 130 days, range 37-451) and time to diagnosis (median in cases thirteen days, range 1-84 days vs median in controls 107 days, range 21-429 days) were significantly less in cases than controls (p=0.04).

**CONCLUSIONS:** rWGS increased the proportion of infants in a regional NICU and PICU who received a timely diagnosis of a genetic disease. Additional, adequately powered studies are needed to determine whether accelerated diagnosis is associated with improved outcomes in this setting. ClinicalTrials.gov Identifier: NCT02225522.

## INTRODUCTION

A premise of pediatric precision medicine is that outcomes are improved by replacement of clinical diagnosis and empiric management with genetic diagnosis and genotype-differentiated treatment^1-9^. The evidence base for pediatric precision medicine is still underdeveloped^10,11^. Ill infants are especially in need of precision medicine since genetic diseases are a leading cause of mortality, particularly in neonatal intensive care units (NICU) and pediatric intensive care units (PICU)^5-7,12-16^. Amongst high-cost health care, NICU treatment is one of the most cost-effective^17-19^. Since disease progression can be very rapid in infants, genetic diagnoses must be made quickly to permit consideration of precision interventions in time to decrease morbidity and mortality^5,6,20-23^. For a few genetic diseases, newborn screening has shown early neonatal diagnosis and rapid, precise intervention to dramatically improve outcomes^24,25^. The potential expansion to newborn diagnosis for symptomatic infants for all 5000 genetic diseases^26^ has been made technically possible by the advent clinical genomic sequencing (whole genome sequencing (WGS) or whole exome sequencing (WES), and next-generation sequencing gene panel tests (NGS)). In particular, rapid WGS (rWGS) can allow genetic diagnosis in two days^20,27^.

There is substantial evidence that a higher proportion of symptomatic children with likely genetic disease receive etiologic diagnoses by WGS and WES than other genetic tests^3-7,6,28-35^. Published NICU or PICU experience with rWGS, however, is limited to case reports and one retrospective study^5,6,20-23^. In the latter, 57% of infants received genetic diagnoses in a median of 23 days (day of life 49)^6^. However, it has not yet been unequivocally demonstrated whether rWGS improves timeliness of genetic diagnosis relative to standard genetic tests. Here we report results of **<u>N</u>**ewborn **<u>S</u>**equencing **<u>I</u>**n **<u>G</u>**enomic medicine and public **<u>H</u>**eal**<u>T</u>**h Randomized Controlled Trial (RCT) 1 (NSIGHT1), the first RCT of genomic testing in patients^24^. Specifically, NSIGHT1 compared rates of genetic diagnosis in NICU and PICU infants with possible genetic diseases at 28 days from enrollment by standard tests alone vs standard tests plus trio rWGS.

## METHODS

Full details of methods are reported in Supplementary Material.

### Trial Design

NSIGHT1 tested the *a priori* hypothesis that rWGS increases the proportion of infants receiving a genetic diagnosis within 28 days in a partially blinded, randomized controlled study in a regional NICU and PICU in a tertiary referral children’s hospital (Children’s Mercy – Kansas City). Enrollment was from October 2014 - June 2016, and follow up until November 2016. Inclusion criteria were infants in the NICU or PICU of age less than four months with illnesses of unknown etiology and one of the following: 1. A genetic test order or genetic consult; 2. A major structural congenital anomaly or at least three minor anomalies; 3. A laboratory test suggested a genetic disease; or 4. An abnormal response to therapy. Exclusion criteria were an existing genetic diagnosis, or features pathognomonic for a chromosomal aberration. The NICU census was reviewed daily for eligible infants by enrollment coordinators. NICU clinicians were notified of eligible infants, who were nominated through a standard form. NICU and PICU clinicians notified families of eligible infants about the study, and enrollment coordinators then approached parents for informed consent. Enrolled infants were randomly assigned to receive standard clinical tests (controls) or standard clinical tests plus trio (infants and parents where available) rWGS (cases; Figure S1). Parents and clinicians were blinded until by day ten, when they were notified of randomization assignment, to allow consideration of crossover to rWGS.

### Rapid Genome Sequencing

rWGS was performed using previously described methods that yielded variant calls within two – seven days of blood draw^5,6,20,27^. Variants were interpreted by board certified molecular geneticists using American College of Medical Genetics guidelines for pathogenic and likely pathogenic classifications^36^. Genotypes were confirmed clinically by Sanger sequencing. Secondary and incidental findings were not reported.

### Standard Genetic Testing

Standard clinical testing for genetic diseases was performed based on clinician judgment, assisted by subspecialist recommendations. The set of genetic tests considered to be standard was developed by two molecular genetics laboratory directors (Supplementary Methods).

### Trial End Points

The primary end point was the diagnostic sensitivity within 28 days of enrollment or first standard test order. Secondary end points were the diagnostic sensitivity by day of life (DOL) 28, total diagnostic sensitivity, time-to-diagnosis, rate of clinical utility (proportion of patients with a change in management related to test results), length of hospitalization, and 6 month mortality rate. Clinical utility was determined by clinician surveys and reviews of the electronic health record by at least two pediatric subspecialist experts in genomic medicine to identify changes in treatments, procedures, consultations, testing, genetic or reproductive counseling, and recommendations for specific follow up related to the diagnosis^37^. A modified Delphi method was used to determine inclusion of change in management where there was disagreement.

### Statistical Analysis

Statistical analyses were based on the intention-to-treat principle. Fisher’s exact test was used to compare 28-day diagnostic rates, total diagnoses, clinical utility of diagnoses, and diagnoses before discharge. A two-sample t-test was performed to compare age at hospital discharge. Kaplan-Meier analyses were used to compare time to diagnosis, which was measured from the date of first standard test order for controls or date of enrollment for cases, and age at diagnosis. Age at death was compared with the log-rank test^38^. When there was evidence of a non-constant hazard ratio, between-group differences were evaluated with the Peto-Peto test^39,40^.

## RESULTS

### Patients

65 (50%) of 129 nominated infants completed the study (Figure 1). 32 infants randomized to rWGS plus standard genetic tests (cases) and 33 to standard tests alone (controls, Figures 1, S1). Phenotypes were highly diverse and typically present at birth (Tables 1, S1). Fewer control infants had cardiovascular findings (6% vs 28%; difference, −22% [95% CI, −40% to −4%]; p=0.02) than cases, which may have affected likelihood for genetic disease (Table 1).

**Figure 1.**
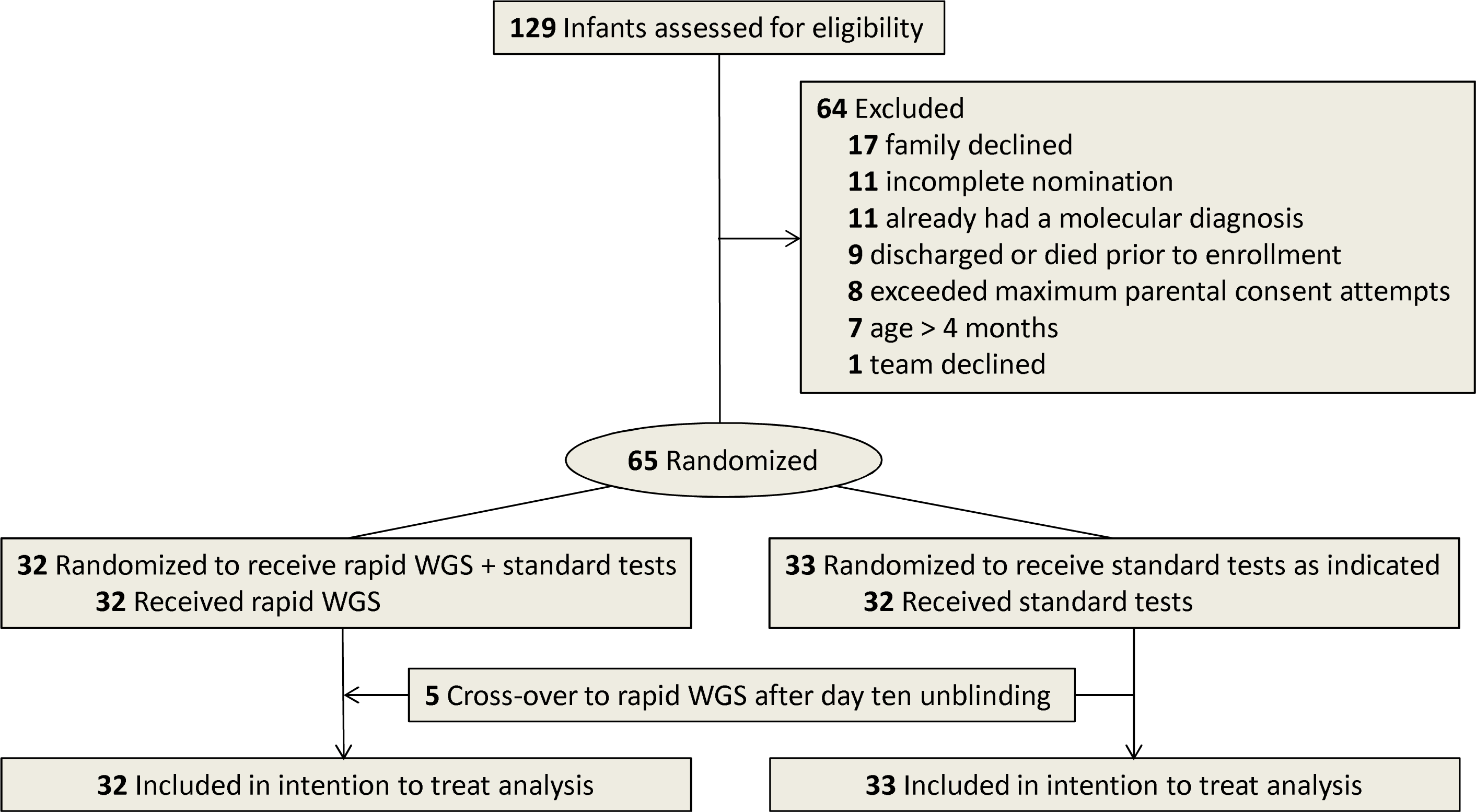
CONSORT flow diagram of NSIGHT1 enrollment and randomization. Major reasons for non-enrollment were family refusal (13%), the infant had a diagnosis that explained the phenotype (9%), and incomplete nominations (9%). At unblinding of clinicians (by 10 days after enrollment), requests were made for compassionate cross-over of 7 (21%) of 33 infants who randomized to standard tests alone to receive rWGS, of which 5 were granted.

**Table 1:** Characteristics of the 65 NSIGHT1 probands.

|  |  | **Cases (rWGS,**<br>**n=32)** | **Controls (n=33)** |
| --- | --- | --- | --- |
| **Sex** | Female (n, %) | 15 (47%) | 11 (33%) |
|  | Male (n, %) | 16 (50%) | 21 (64%) |
|  | Undetermined (n, %) | 1 (3%) | 1 (3%) |
| **Demographics** | Caucasian (n, %) | 25 (78%) | 27 (82%) |
|  | African, African American (n, %) | 2 (6%) | 1 (3%) |
|  | Other Race (n, %) | 5 (16%) | 5 (15%) |
|  | Hispanic (n, %) | 2 (6%) | 3 (9%) |
|  | Consanguinity (n, %) | 1 (3%) | 2 (6%) |
| **Birth Characteristics** | Gestational Age (Average, wks) | 36.0 | 35.9 |
|  | Weight (average, kg) | 2.5 | 2.4 |
|  | Low Birth Weight (<2500 g, n, %) | 14 (44%) | 9 (27%) |
|  | Extremely Low Birth Weight (<1000 g, n, %) | 1 (3%) | 3 (9%) |
|  | APGAR at 1 minute (Average) | 6.1 | 5.1 |
|  | APGAR at 5 minutes (Average) | 7.8 | 6.4 |
|  | Symptom Onset (Average day of life) | 2.3 | 2.1 |
| **Primary System Involved by Disease** | Congenital Anomalies/Musculoskeletal | 10 (31%) | 13 (39%) |
|  | Neurological | 5 (16%) | 11 (33%) |
|  | Cardiovascular Findings | 9 (28%) | 2 (6%) |
|  | Endocrine/Metabolic | 1 (3%) | 3 (9%) |
|  | Respiratory Findings | 4 (13%) | 0 (0%) |
|  | Other | 2 (6%) | 2 (6%) |
|  | Renal | 1 (3%) | 2 (6%) |
|  | Dermatologic | 1 (3%) | 0 (0%) |
|  | Multiple System | 1 (3%) | 0 (0%) |
|  | Hepatic | 0 (0%) | 0 (0%) |
| **Enrollment & Standard Clinical Tests** | Day of life at Enrollment (Average, range) | 22.8 (1-101) | 22.0 (1-80) |
|  | Subjects receiving Standard Clinical Tests (n, %) | 30 (93.8%) | 33 (100%) |
|  | Day of Life 1<sup>st</sup> Standard Test Ordered (Average, range) | 11.6 (0-66) | 15.6 (0-120) |
|  | Standard Tests Ordered (Average, range) | 2.8 (0-7) | 3.4 (1-10) |
|  | Probands receiving Standard Clinical NGS Tests (n, %) | 14 (44%) | 21 (64%) |
|  | Standard NGS Tests Ordered (n, % of total Standard Tests) | 14 (16%) | 29 (26%) |
|  | Diagnosis by Standard Test (n, %) | 7 (22%) | 8 (24%) |
|  | DOL Diagnosis by Standard Clinical Test (median, range) | 66 (16-151) | 130 (37-451) |
|  | Time to Diagnosis by Standard Clinical Test (Average, range) | 45 (16-150) | 110 (31-450) |

### Standard Diagnostic Tests

The proportion of infants receiving standard genetic tests and age at first standard test order were similar in both arms (Table 1). Infants received an average of 3.1 (range 0-10) standard genetic tests (Table 1, S3). 21 (64%) of 33 control infants received non-expedited NGS, WES or WGS standard tests, compared with fourteen (44%) of 32 cases (Table 1, S3). The average age at first standard test order was 14 days (range 0–120 days). Standard tests yielded fifteen (43%) genetic diagnoses in the 35 subjects tested, seven (50%) in 14 cases, and eight (38%) in 21 controls (Table 2, S4). Of note, five (8%) diagnoses by standard tests were not detected by rWGS at the time of study: Four (6%) were copy number or structural variants and one (2%) was a change in DNA methylation. The median time from first standard test order to diagnosis was 64 days (range 16-450 days). The average age at diagnosis by standard genetic tests was 113 days (range 16– 451 days). Six (9.4%) of 64 infants received a diagnosis by standard tests prior to hospital discharge (Table S5).

**Table 2:** Presentations and characteristics of the twenty one infants who received diagnoses.

| Patient ID | Study Arm | Dx Type | Mode of Dx | Primary Clinical Features<sup>1</sup> | Diagnosis | Gene | Inheritance Pattern | De novo or inherited | Variant Chromosomal (Chr)<sup>2</sup> or Gene (c.) Coordinates | Variant Protein Coordinates |
| --- | --- | --- | --- | --- | --- | --- | --- | --- | --- | --- |
| 5004 | Case | Partial | Std | Cleft palate micrognathia hypoglycemia hyperinsulinemia thrombocytopenia | Chr 7p duplication syndrome | *n.a.* | n.d. | n.d. | Gain 7p22.3-p15.2 Chr7:43360-26463160dup | n.a. |
| 5007 | Control | Full | WGS & Std | Polymicrogyria Intractable seizures Epileptic encephalopathy | Congenital disorder of glycosylation type Ik | *ALG1* | Autosomal Recessive | Inherited | c.15C>A and c.149A>G | p.C5* + p.Q50R |
| 5008 | Case | Full | Std | Complete atrioventricular canal defect Hypospadias IUGR Dysmorphic features | Chr 8p23 deletion syndrome | *n.a.* | n.d. | n.d. | Chr8:158048-6999114del 10054927-10479436dup 10479473-11882401del | n.a. |
| 5011 | Control | Full | Std | Hypotonia Cryptorchidism Aniridia | XL myotubular myopathy-1 Aniridia | *MTM1*<br>*PAX6* | X-Linked Recessive; Autosomal Dominant | n.d.<br>Inherited | c.137-3T>G;<br>c.1268A>T | n.a.<br>p.*423L |
| 5014 | Control | Full | Std | Hyperglycemia | Transient neonatal diabetes | *ZFP57* | n.d. | n.a. | Hypomethylation 6q24 | n.a. |
| 5023 | Case | Full | WGS & Std | Hyponatremia SGA/IUGR Pseudohypoadosteronism | Pseudohypoadosteronism type I | *NR3C2* | Autosomal Dominant | Inherited | c.1951C>T | p.R651* |
| 5025 | Control | Full | Std | Micrognathia Cleft palate Abnormal facies Right thumb hypoplasia | Nager type acrofacial dysostosis | *SF3B4* | Autosomal Dominant | *de novo*; Inherited | c.1088-3C>G; c.1058C>A | n.a.; p.P353H |
| 5026 | Control | Full | Std | Hirsutism Mild Synophrys Mild Micrognathia Camptodactyly Renal cysts | Cornelia de Lange syndrome 1 | *NIPBL* | Autosomal Dominant | *de novo* | c.5057del | p.L1686Rfs*7 |
| 5027 | Control | Full | Std | IUGR Cleft palate Micrognathia Skin tags Poor gag reflex | Chr 1p36 deletion syndrome | *n.a.* | n.d. | n.d. | Loss arr 1p36.11 Chr1:24100645-25003678del | n.a. |
| 5030 | Case | Full | Std | Seizures Poor feeding | AD Nocturnal Frontal Lobe Epilepsy | *CHRNA4* | Autosomal Dominant | *de novo* | Heterozygous deletion of the entire CHRNA4 gene | n.a. |
| 5035 | Case | Full | WGS | Microcephaly | Primary AR Microcephaly 5 | *ASPM* | Autosomal Recessive | Inherited | c.3428dupT;<br>c.8191_8192del | p.L1144Vfs*16;<br>p.E2731Kfs*19 |
| 5036 | Case | Full | WGS | Central apnea | Congenital Central Hypoventilation Syndrome | *PHOX2B* | Autosomal Dominant | *de novo* | PHOX2B ALA EXP | p.A260(9) |
| 5038 | Case | Full | WGS | Situs inversus | Primary Ciliary Dyskinesia type 7 | *DNAH11* | Autosomal Recessive | Inherited | c.6244C>T; c.6776A>T and c.8567T>C | p.R2082*;<br>p.D2259V and p.V2856A |
| 5042 | Case | Full | WGS | Profound hypotonia Respiratory distress Myoclonic jerks | AD Mental Retardation 31 | *PURA* | Autosomal Dominant | *de novo* | c.458_459dupC | p.K154Qfs*47 |
| 5048 | Case | Full | WGS | Seizures | Early Infantile Epileptic Encephalopathy 14 | *KCNT1* | Autosomal Dominant | *de novo* | c.1420C>T | p.R474C |
| 5051 | Case | Full | WGS | Perinatal ascites; cholestasis | Dehydrated Hereditary Stomatocytosis | *PIEZO1* | Autosomal Dominant | Inherited | c.6058G>A | p.A2020T |
| 5053 | Control | Full | WGS & Std | Altered mental status Decreased deep reflexes Hypotonia Cryptorchidism | XL Myotubular Myopathy | *MTM1* | X-linked Recessive | *de novo* | c.567_569delTAA | p.N189del |
| 5057 | Case | Full | WGS & Std | Dysmorphic features Cardiac anomalies Failed hearing screen | Noonan Syndrome | *SOS1* | Autosomal Dominant | *de novo* | c.2536G>A | p.E846K |
| 5059 | Case | Full | WGS & Std | HLHS Hydrocephalus Multiple congenital anomalies | Coffin-Siris Syndrome | *ARID1A* | Autosomal Dominant | *de novo* | c.1207C>T | p.Q403* |
| 5061 | Case | Partial | WGS & Std | Hypotonia Absent gag reflex Exaggerated startle reflex | Hyperekplexia | *GLRA1* | Autosomal Dominant | *de novo* | c.373G>A | p.D125N |
| 5062 | Control | Full | Std | Bicuspid aortic valve, Hypotonia, Leukocytosis | Central Core Disease of Muscle | *RYR1* | Autosomal Dominant | *de novo* | c.14581C>T | p.R4861C |

<sup>1</sup>Full clinical features are shown in Table S1; <sup>2</sup>GRCh37; Chr: Chromosome; Std: standard genetic test.

### Rapid Whole Genome Sequencing

Ten of 32 cases (31%) received diagnoses by rWGS (Table 2, Table S4). Including five crossovers, 12 (32%) of 37 infants received rWGS diagnoses (Table 2, S5). On average, enrollment occurred on DOL 22 (range one −101; Table 1), an average of eight days later than standard tests. The median time to rWGS diagnosis, including clinical confirmatory testing, was fourteen days (range eight – 35 days; Table S5). The median age at WGS diagnosis in patients randomized to rWGS was 28.5 days (range 14 – 90 days). Among crossovers, the median age at WGS diagnosis was 94.5 days.

### Diagnoses

Twenty-two genetic diagnoses were reported in 21 (32%) of 65 infants (Table 2). The most common mechanism was *de novo* variant occurrence (eleven of eighteen (61%) diagnoses; Table 2). The most common inheritance pattern was autosomal dominant (thirteen of eighteen (72%) diagnoses). Cross-over to rWGS was requested for seven (21%) of the 33 controls. Five were granted, yielding two diagnoses. In both, diagnosis by rWGS occurred first but was recapitulated by standard tests (Table 2). Twenty (31%) of the 65 infants (91% of those with a diagnoses) had attendant changes in management (Table S4).

### Early study termination

The study was terminated after 21 months due to growing availability of NGS panels, WES and WGS as standard tests, which shifted the baseline of comparison over the course of the study. These were associated with high rates of cross-over requests and higher utilization of NGS panel, WES or WGS standard genetic tests among controls (64% including cross overs) than cases (44%; Table S3).

### End-Point Testing

End-points were analyzed on the basis of intention to treat (Figures 1, S1). The primary end point, rate of genetic diagnosis within 28 days of enrollment, was higher in cases (31%, ten of 32) than controls (3%, one of 33; difference, 28% [95% CI, 10% to 46%]; p=0.003 Table 3, Figure 2). For neonates enrolled within the first 25 days of life, the rate of diagnosis by DOL 28 was higher in cases (32%, seven of 22) than controls (0%, zero of 33; difference, 32% [95% CI, 11% to 53%]; p<0.01; Table 3). Age at diagnosis and time to diagnosis differed significantly between arms, after accounting for non-proportional rates of diagnosis (Table 4, Table S5): The median age at diagnosis in cases was 25 days (range 14-90 days) vs median in controls was 130 days (range 37-451). The median time to diagnosis in cases was 13 days (range 1-84 days) vs median in controls 107 days (range 21-429 days).

**Table 3:** Comparison of Primary and Secondary End-Points.

|  | rWGS + Standard Testing | Standard Testing (Including crossovers) | P-Value | Statistical Test |
| --- | --- | --- | --- | --- |
| **Number of subjects** | 32 | 33 |  |  |
| **Primary End-Point** |  |  |  |  |
| Diagnosis within 28 days of standard test order/enrollment (n, %) | 10 (31%) | 1 (3%) | 0.003<sup>1</sup> | Fisher's exact test |
| **Secondary End-Points** |  |  |  |  |
| Diagnosis by DOL 28 (n, %) | 7 (32%) | 0 (0%) | 0.004<sup>1</sup> | Fisher's exact test |
| Total Diagnoses (n, %) | 13 (41%) | 8 (24%) | 0.19 | Fisher's exact test |
| Clinical Utility of Diagnoses (n, %) | 13 (41%) | 7 (21%) | 0.11 | Fisher's exact test |
| DOL Hospital Discharge (average, range) | 66.3 (3-456) | 68.5 (4-341) | 0.91 | Two sample t-test |
| Diagnosis before Discharge (n, %) | 9 (28%) | 3 (9%) | 0.06 | Fisher's exact test |
| Mortality at 180 days (n, %) | 4 (13%) | 4 (12%) | n.d. |  |
| Age at death (days; median, range) | 62 (14-228) | 173 (4-341) | 0.93 | Log rank test |

<sup>1</sup>Fisher's exact test p-value both for all patients and in a sensitivity analysis, in which patients with a partial diagnosis (5004 and 5061) were considered undiagnosed. DOL: day of life.

**Figure 2.**
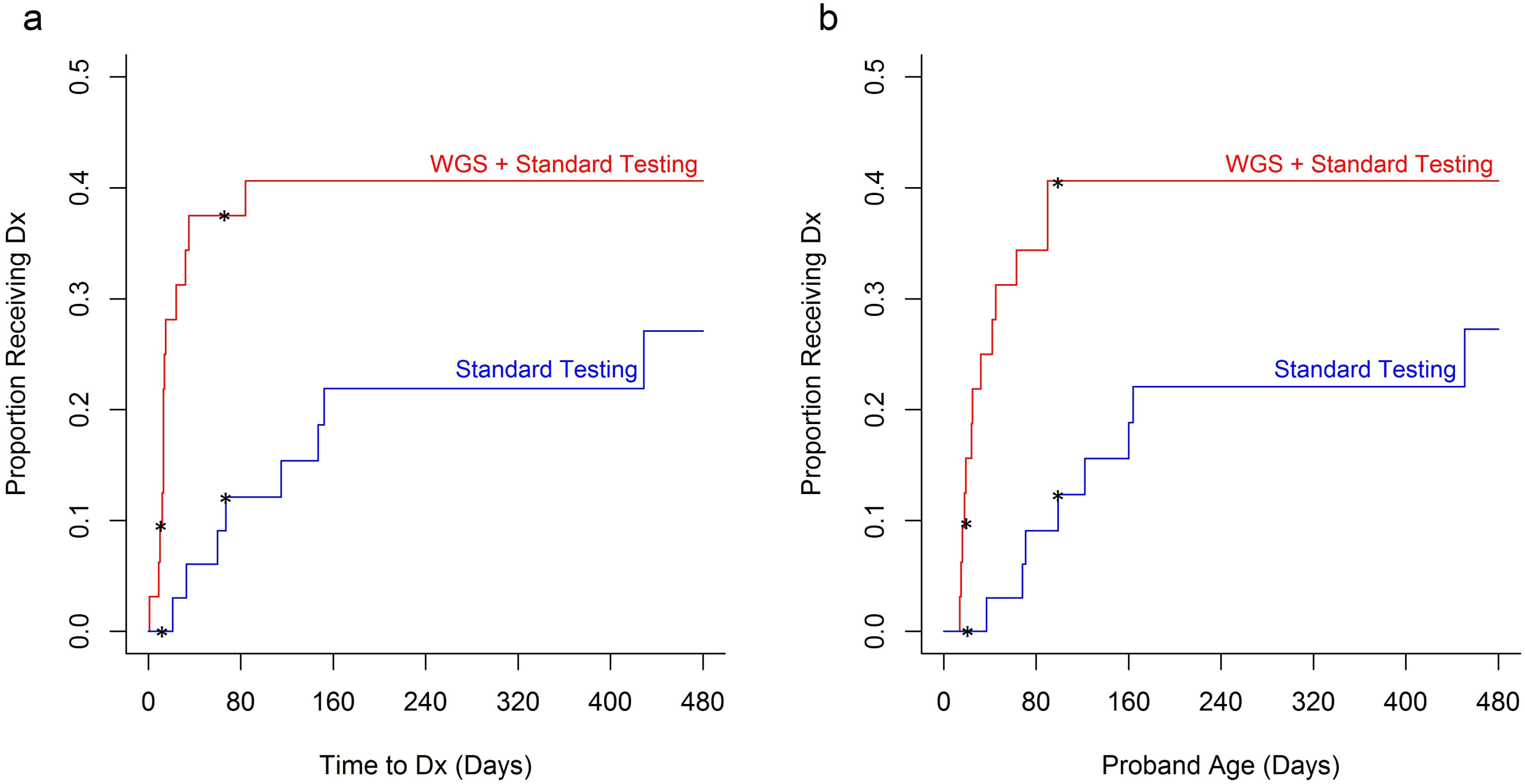
Kaplan-Meier curves of time to diagnosis in cases and controls. The cumulative probability of a diagnosis (Dx) in cases (infants randomized to receive rWGS plus standard genetic tests; shown in red; n=32) and controls (infants randomized to standard genetic tests alone; shown in blue; n=33). Differences in probability of receiving a diagnosis were significant between the two arms from day 12 – 67 after enrollment (panel a, asterisks) and DOL 19 - 99 (panel b, asterisks).

Five secondary end-points did not differ significantly between arms (Table 3, 4, S4). They were the proportion of infants in whom diagnoses had clinical utility (41% of cases vs 21% of controls; difference, 19% [95% CI, −3% to 42%]), proportion of infants with a change in medical management (clinical utility, 22% of cases vs 9% of controls; difference, 13% [95% CI, −5% to 30%]), proportion of patients who received diagnoses prior to hospital discharge (28% of cases vs 9% of controls; difference, 19% [95% CI, 0% to 38%]), average length of NICU/PICU stay, 6-month mortality, and age at death.

**Table 4:** Comparison of age at diagnosis and time to diagnosis between cases (rWGS plus standard tests) and controls (standard tests alone).

|  | Original analysis<sup>1</sup> |  | Sensitivity analysis<sup>2</sup> |  |
| --- | --- | --- | --- | --- |
|  | p-value for non-proportional hazards | p-value for a difference in overall Dx rates | p-value for non-proportional hazards | p-value for a difference in overall Dx rates |
| Age at Diagnosis | 0.002 | 0.043 | 0.003 | 0.15 |
| Time to diagnosis from enrollment/1st test ordered (depending on which was earliest for cases) and from 1<sup>st</sup> test ordered (controls) | 0.002 | 0.040 | 0.002 | 0.11 |

<sup>1</sup>Peto-Peto test used instead of log-rank test due to evidence of non-proportional hazards; <sup>2</sup>Peto-Peto test when patients with a partial Dx (5004 and 5061) considered undiagnosed.

## Discussion

NICU and PICU infants receiving trio rWGS plus standard clinical testing had a higher rate of genetic diagnosis and shorter time to diagnosis than infants receiving standard tests alone. In intention to treat analysis, rWGS was associated with significantly more genetic diagnoses within 28 days of enrollment (31%, 10 of 32) than standard tests alone (3%, 1 of 33; difference, 28% [95% CI, 10% to 46%]; p=0.003). The rate of neonatal (DOL 28) diagnosis was higher in cases (32%, 7 of 22) than controls (0%, 0 of 23; difference, 32% [95% CI, 11% to 53%]; p=0.004). Of note, standard genetic testing was ordered an average of 8 days before enrollment, which benefitted the control arm over rWGS cases for these analyses. Nevertheless, age at diagnosis and time to diagnosis were significantly shorter in rWGS cases, after accounting for non-proportional rates of diagnosis.

The rate of genetic diagnosis by rWGS in a NICU or PICU was reported previously in one cohort^6^. Enrollment in that study was at average DOL 26 (vs DOL 22 herein). The rate of diagnosis by rWGS therein was 14% (5 of 35) by DOL 28, and 34% (12 of 35) within 28 days of enrollment, which were similar to herein (32% and 31%, respectively). The total rate of genetic diagnosis by rWGS herein (32%) was within the range reported for WGS and WES studies^3-7,6,28-35^.

Timely return of rWGS diagnoses was limited by two research factors that may not be part of routine clinical practice: firstly, confirmatory testing by “the clinically accepted standard” was required for research rWGS diagnoses – but is not necessarily required for laboratory developed NGS tests – which lengthened the time to rWGS diagnosis by 7 – 10 days. Indeed, all diagnostic rWGS findings in the current study were concordant with orthologous methods. For well covered, pathogenic and likely pathogenic, single nucleotide variants in regions of high WGS quality, a median time-to-result of five days is anticipated^6,20,27^.

Secondly, enrollment occurred relatively late during the NICU or PICU stay (DOL 22). While parents are interested in receipt of genomic sequencing at birth, an enrollment rate of 6% was reported for WES in NICU infants in another cohort^41,42^. Delay in enrollment herein reflected two logistical factors. First, since a criterion for enrollment was suspicion by the provider of an underlying genetic disease, nomination was often delayed until a genetic test or consult had been ordered. In such cases, the time of enrollment delayed the study test, rWGS, compared to standard testing; nevertheless, there was still a decreased time to diagnosis with rWGS. Secondly, NSIGHT1 required informed consent from both parents; the logistics and complexity of obtaining informed consent in a NICU or PICU setting are arduous. In future studies, it will be important to seek enrollment close to day of admission. This would be facilitated by simpler enrollment criteria, requirement of informed consent from a single parent, and limiting eligibility for enrollment to within several days of admission.

Clinical WGS continues to improve with respect to rate of genetic diagnosis and time to diagnosis^27^. In particular, the diagnostic rate is increasing through ongoing identification of novel disease genes, improved reference genome sequences, and better identification of disease-causing copy number, repeat expansion, regulatory, splicing and structural variations^32,43-50^. These recent advances were not reflected in the current study. WES and WGS have similar analytic performance for exonic and splicing variants, which comprised seventeen of twenty two diagnoses. However, four diagnoses were associated with copy number or structural variants, for which WGS has superior analytic performance to WES. rWGS is methodologically simpler than WES, and thus two days faster than possible with rapid WES.

NSIGHT1 was terminated early, primarily due to loss of equipoise noted by some nominating clinicians during the study. Some practitioners grew to regard randomization to standard tests alone to be an inferior intervention than standard tests plus trio rWGS. This was associated with seven (21% of controls) requests to cross-over control infants to the rWGS arm following clinician un-blinding, five of which were granted. It was also associated with a higher rate of order of NGS panel, WES or WGS standard genetic tests in controls (64%) than cases (44%). Standard genomic sequencing tests accounted for 63% (5) of the 8 genetic diagnoses in controls. As a result, there was not a significant difference between arms in the total number of genetic diagnoses, a secondary end-point (41% [13] diagnoses among 32 infants in the rWGS arm, 24% [8] of 33 in controls; difference, 16% [95% CI, −6% to 39%]; p=0.19). Future pragmatic RCT designs in genomic medicine will require careful attention to the principle of equipoise and to the rapid evolution of clinical NGS-based testing^51-52^. The more widespread use of gene panel testing in the NICU during the course of this study was a significant departure from our experience at study conception. Our study was not intended to evaluate the relative diagnostic yield of panel testing over rWGS. Consequently, the study was not powered to evaluate the non-inferiority of panels over rWGS.

The rationale for rWGS in NICU infants is to enable consideration of acute precision interventions in time to decrease morbidity and mortality^5,6,21-24^. In two prior studies of genomic sequencing in infants, genetic diagnoses led to precision medicine that was considered life-saving in 5%, and that avoided major morbidity in 6% ^6,7^. In those studies, early diagnosis (DOL 49) led to greater implementation of precision medicine (65%) than later diagnosis (DOL 374, 39%), particularly with regard to palliative care guidance. As in the current study, assessments of clinical utility were based on actual changes in management, which were limited by clinician experience with genomic medicine and rare genetic diseases. This is a major challenge for NICU and PICU implementation of genomic medicine for rare genetic diseases^53^. Unfortunately, early termination of the current study resulted in loss in power for the secondary endpoints: There were not significant differences in the overall rate of clinical utility of diagnoses, length of admission, rate of diagnosis before discharge, mortality and age at death. The clinical utility of diagnoses and rate of diagnosis before hospital discharge trended towards being higher in the rWGS arm (difference, 19% [95% CI, −3% to 42%], p=0.11, and 19% [95% CI, 0% to 38%], p=0.06, respectively). Additional studies are clarify whether shorter time to diagnosis is associated with changes in clinical utility of diagnoses, outcomes, or healthcare utilization.

## Conclusions

Among infants with suspected genetic diseases in a regional NICU or PICU, the addition of rWGS decreased the time to diagnosis. We suggest that rWGS should be considered as a first-tier genetic test in NICU and PICU infants with suspected genetic diseases^7^. Since genetic diseases are among the leading cause of death in the NICU and PICU, as well as overall infant mortality, implementation of rWGS is likely to have broad implications for the practice of neonatalology.

## Article Information

### Author contributions

Concept and design: SFK, LKW, JEP.

Acquisition, analysis or interpretation of data: All authors.

Drafting of manuscript: SFK, JEP, MMC, NS, JAC.

Critical Revision of the Manuscript for Important Intellectual Content: DPD, PMA, SLC, MAC, RTF, JAH, HK, RJM, JLR, SLT.

Statistical analysis: Clark.

Obtained Funding: Kingsmore, Willig, Leeder.

Administrative, Technical or Material Support: SFK, JAC.

Study Supervision: JEP, SFK, JAC.

### Additional Contributions

We thank Drs. John Lantos, Sarah Soden and Howard Kilbride for advice and assistance. *A Deo lumen, ab amicis auxilium*.

### Data and material availability

Data are available at LPDR (https://www.nbstrn.org/research-tools/longitudinal-pediatric-data-resource).

## REFERENCES

1. Green ED, Guyer MS, National Human Genome Research I. Charting a course for genomic medicine from base pairs to bedside. Nature. 2011; 470(7333):204–13.

2. Worthey EA, Mayer AN, Syverson GD, et al. Making a definitive diagnosis: successful clinical application of whole exome sequencing in a child with intractable inflammatory bowel disease. Genet Med. 2011; 13(3):255–62.

3. Bainbridge MN, Wiszniewski W, Murdock DR, et al. Whole-genome sequencing for optimized patient management. Sci Transl Med. 2011; 3(87): 87re3.

4. Dixon-Salazar TJ, Silhavy JL, Udpa N, et al. Exome sequencing can improve diagnosis and alter patient management. Sci Transl Med. 2012; 4(138):138ra78.

5. Soden S, Saunders CJ, Willig LK, et al. Effectiveness of exome and genome sequencing guided by acuity of illness for diagnosis of neurodevelopmental disorders. Sci Transl Med. 2014; 6(265):265ra168.

6. Willig LK, Petrikin JE, Smith LD, et al. Whole-genome sequencing for identification of Mendelian disorders in critically ill infants: a retrospective analysis of diagnostic and clinical findings. Lancet Respir Med. 2015; 3:377–87.

7. Stark Z, Tan TY, Chong B, et al. A prospective evaluation of whole-exome sequencing as a first-tier molecular test in infants with suspected monogenic disorders. Genet Med. 2016;18:1090–6

8. Manolio TA, Abramowicz M, Al-Mulla F, et al. Global implementation of genomic medicine: We are not alone. Sci Transl Med. 2015; 7(290):290ps13.

9. Collins FS, Varmus H. A new initiative on precision medicine. N Engl J Med. 2015; 372(9):793–5.

10. Phillips KA, Deverka PA, Sox HC, et al. Making genomic medicine evidence-based and patient-centered: a structured review and landscape analysis of comparative effectiveness research. Genet Med. 2017 Apr 13.

11. Roberts MC, Kennedy AE, Chambers DA, et al. The current state of implementation science in genomic medicine: opportunities for improvement. Genet Med. 2017 Jan 12.

12. Xu J, Murphy SL, Kochanek KD, Arias E. Mortality in the United States, 2015. NCHS Data Brief. 2016;267:1–8.

13. Wilkinson DJ, Fitzsimons JJ, Dargaville PA, et al. Death in the neonatal intensive care unit: changing patterns of end of life care over two decades. Arch Dis Child Fetal Neonatal Ed. 2006; 91(4):F268–71.

14. Hagen CM, Hansen TW. Deaths in a neonatal intensive care unit: a 10-year perspective. Pediatr Crit Care Med. 2004; 5(5):463–68.

15. O’Malley M, Hutcheon RG. Genetic disorders and congenital malformations in pediatric long-term care. J Am Med Dir Assoc. 2007; 8(5):332–34.

16. Stevenson DA, Carey JC. Contribution of malformations and genetic disorders to mortality in a children’s hospital. Am J Med Genet A. 2004;126A(4):393–97.

17. Lantos JD, Meadow WL. Costs and end-of-life care in the NICU: lessons for the MICU? J Law Med Ethics. 2011; 39(2):194–200.

18. Couce ML, Bana A, Boveda MD, et al. Inborn errors of metabolism in a neonatology unit: impact and long-term results. Pediatr Int. 2011; 53(1):13–17.

19. Weiner J, Sharma J, Lantos J, Kilbride H. How infants die in the neonatal intensive care unit: trends from 1999 through 2008. Arch Pediatr Adolesc Med. 2011; 165(7):630–34.

20. Saunders CJ, Miller NA, Soden SE, et al. Rapid whole-genome sequencing for genetic disease diagnosis in neonatal intensive care units. Sci Transl Med. 2012; 4(154):154ra135.

21. Priest JR, Ceresnak SR, Dewey FE, et al. Molecular diagnosis of long-QT syndrome at 10 days of life by rapid whole genome sequencing. Heart Rhythm. 2014; 11(10):1707–13.

22. Farnaes L, Nahas SA, Chowdhury S, et al. Rapid whole genome sequencing identifies a novel GABRA1 variant associated with West syndrome. Cold Spring Harb Mol Case Stud. 2017 May 11.

23. Hildreth A, Wigby K, Chowdhury S, et al. Rapid whole genome sequencing identifies a novel homozygous NPC1 variant associated with Niemann-Pick Type C1 Disease in a 7 week old male with cholestasis. Cold Spring Harb Mol Case Stud. 2017 May 26.

24. Berg JS, Agrawal PB, Bailey DB Jr, et al. Newborn Sequencing in Genomic Medicine and Public Health. Pediatrics. 2017; 139(2)pii: e20162252.

25. Downing GJ, Zuckerman AE, Coon C, Lloyd-Puryear MA. Enhancing the quality and efficiency of newborn screening programs through the use of health information technology. Semin Perinatol. 2010; 34(2):156–62.

26. Online Mendelian Inheritance in Man, OMIM. (Accessed on August 4, 2017, at http://www.omim.org/ http://www.omim.org/).

27. Miller NA, Farrow EG, Gibson M, et al. A 26-hour system of highly sensitive whole genome sequencing for emergency management of genetic diseases. Genome Med. 2015;7:100.

28. Vissers LE, van Nimwegen KJ, Schieving JH, et al. A clinical utility study of exome sequencing versus conventional genetic testing in pediatric neurology. Genet Med. 2017 Mar 23.

29. Stavropoulos DJ, et al. Whole-genome sequencing expands diagnostic utility and improves clinical management in paediatric medicine. npj Genomic Medicine. 2016 1:15012.

30. Tarailo-Graovac M, et al. Exome Sequencing and the Management of Neurometabolic Disorders. N Engl J Med. 2016 374(23):2246–55.

31. Baldridge D, et al. The Exome Clinic and the role of medical genetics expertise in the interpretation of exome sequencing results. Genet Med. 2017 Mar 2.

32. Ellingford JM, Barton S, Bhaskar S, et al. Whole Genome Sequencing Increases Molecular Diagnostic Yield Compared with Current Diagnostic Testing for Inherited Retinal Disease. Ophthalmology. 2016 123(5):1143–50.

33. Farwell KD, Shahmirzadi L, El-Khechen D, et al. Enhanced utility of family-centered diagnostic exome sequencing with inheritance model-based analysis: results from 500 unselected families with undiagnosed genetic conditions. Genet Med. 2015 Jul;17(7):578–86.

34. Taylor JC, et al. Factors influencing success of clinical genome sequencing across a broad spectrum of disorders. Nat Genet. 2015; 47(7):717–26.

35. Valencia CA, Husami A, Holle J, et al. Clinical Impact and Cost-Effectiveness of Whole Exome Sequencing as a Diagnostic Tool: A Pediatric Center's Experience. Front Pediatr. 2015;3:67.

36. Richards S, Aziz N, Bale S, et al. Standards and guidelines for the interpretation of sequence variants: a joint consensus recommendation of the American College of Medical Genetics and Genomics and the Association for Molecular Pathology. Genet Med. 2015; 17(5):405–24

37. ACMG Board of Directors. Clinical utility of genetic and genomic services: a position statement of the American College of Medical Genetics and Genomics. Genet Med. 2015; 17(6):505–7.

38. Grambsch PM, Therneau TM. Proportional hazards tests and diagnostics based on weighted residuals. Biometrika. 1994;81:515–26.

39. Peto R, Peto J. Asymptotically efficient rank invariant test procedures. J R Stat Soc Ser A. 1972;135:185–207.

40. Leton E, Zuluaga P. Equivalence between score and weighted tests for survival curves. Commun Stat Theory Methods. 2001;30:591–608.

41. Green RC, Holm IA, Rehm HL, et al. The BabySeq Project: Preliminary findings from a randomized trial of exome sequencing in newborns. Presented at the American Society of Human Genetics 2016 Annual Meeting. Vancouver, B.C., Canada. Oct 19, 2016. https://ep70.eventpilot.us/web/page.php?page=IntHtml&project=ASHG16&id=160122602

42. Waisbren SE, Bäck DK, Liu C, et al. Parents are interested in newborn genomic testing during the early postpartum period. Genet Med. 2015; 17(6):501–504.

43. Eldomery MK, Coban-Akdemir Z, Harel T, et al. Lessons learned from additional research analyses of unsolved clinical exome cases. Genome Med. 2017 9(1):26.

44. Känsäkoski J, Jääskeläinen J, Jääskeläinen T, et al. Complete androgen insensitivity syndrome caused by deep intronic pseudoexon-activating mutation in the androgen receptor gene. Sci Rep. 2016 6:32819.

45. Hartmannová H, Piherová L, Tauchmannová K, et al. Acadian variant of Fanconi syndrome is caused by mitochondrial respiratory chain complex I deficiency due to a non-coding mutation in complex I assembly factor NDUFAF6. Hum Mol Genet. 2016 25(18):4062–4079.

46. Smedley D, Schubach M, Jacobsen JO, et al. A Whole-Genome Analysis Framework for Effective Identification of Pathogenic Regulatory Variants in Mendelian Disease. Am J Hum Genet. 2016 Sep 1; 99(3):595–606.

47. Noll AC, Miller NA, Smith LD, et al. Clinical detection of deletion structural variants in whole-genome sequences. npj Genomic Medicine. 2016;1:16026. doi:10.1038/npjgenmed.2016.26

48. Schneider VA, Graves-Lindsay T, Howe K, et al. Evaluation of GRCh38 and de novo haploid genome assemblies demonstrates the enduring quality of the reference assembly. Genome Res. 2017 27(5): 849–864.

49. Kojima K, Kawai Y, Misawa K, et al. STR-realigner: a realignment method for short tandem repeat regions. BMC Genomics. 2016; 17(1):991.

50. Dolzhenko E, van Vugt J.J.F.A, Shaw RJ, et al. Detection of long repeat expansions from PCR-free whole-genome sequence data. bioRxiv July 20, 2017. doi: https://doi.org/10.1101/093831

51. London AJ. Equipoise in Research: Integrating Ethics and Science in Human Research. JAMA. 2017; 317(5):525–526.

52. Lilford RJ, Jackson J. Equipoise and the ethics of randomization. J R Soc Med. 1995; 88(10):552–9.

53. Bowdin S, Gilbert A, Bedoukian E et al: Recommendations for the integration of genomics into clinical practice. Genet Med. 2016; 18(11): 1075–1084.

54. Gaff CL, Winship IM, Forrest SM, et al. Preparing for genomic medicine: a real world demonstration of health system change. npj Genomic Medicine 2017; 2: 16. doi:10.1038/s41525–017-0017–4

